# Cross-linking mass spectrometric analysis of the endogenous TREX complex from *S. cerevisiae*

**DOI:** 10.1101/2023.06.23.546279

**Authors:** Carina Kern, Christin Radon, Wolfgang Wende, Alexander Leitner, Katja Sträßer

## Abstract

The conserved TREX complex has multiple functions in gene expression such as transcription elongation, 3’ end processing, mRNP assembly and nuclear mRNA export as well as the maintenance of genomic stability. In *S. cerevisiae*, TREX is composed of the pentameric THO complex, the DEAD-box RNA helicase Sub2, the nuclear mRNA export adaptor Yra1 and the SR-like proteins Gbp2 and Hrb1. Here, we present the structural analysis of the endogenous TREX complex of *S. cerevisiae* purified from its native environment. To this end, we used cross-linking mass spectrometry to gain structural information on regions of the complex that are not accessible to classical structural biology techniques. We also used negative-stain electron microscopy to investigate the organization of the cross-linked complex used for XL-MS by comparing our endogenous TREX complex with recently published structural models of recombinant THO-Sub2 complexes. According to our analysis, the endogenous yeast TREX complex preferentially assembles into a dimer. The overall structures of the recombinant yeast THO-Sub2 complexes strongly resemble the structural conformation of the monomers and the dimer interface of the endogenous TREX complex.

## INTRODUCTION

In eukaryotes, nuclear mRNP assembly is a key step of gene expression. Processing of the nascent mRNA occurs as it is synthesized by RNA polymerase II (RNAPII) and includes capping, splicing and 3’ end formation. In parallel, the mRNA associates with multiple nuclear RNA binding proteins (RBPs) that package it into a messenger ribonucleoprotein particle (mRNP) (Meinel and Strasser 2015; Singh et al. 2015; Wegener and Muller-McNicoll 2019; Wende et al. 2019; Khong and Parker 2020). Correct mRNP assembly is mutually dependent on mRNA processing and a prerequisite for nuclear mRNA export. Furthermore, the composition of mRNPs can also affect later stages of the mRNA life cycle such as mRNA localization, translation and degradation in the cytoplasm (Gehring et al. 2017).

A key player in nuclear mRNP assembly is the evolutionarily conserved TREX complex, which couples <u>tr</u>anscription to nuclear mRNA <u>ex</u>port (Jimeno et al. 2002; Strasser et al. 2002; Zenklusen et al. 2002). In *S. cerevisiae*, the TREX complex is composed of the pentameric THO complex (Tho2, Hpr1, Mft1, Thp2 and Tex1), the ATP-dependent DEAD-box RNA helicase Sub2, the nuclear mRNA export adaptor Yra1 and the SR-like proteins Gpb2 and Hrb1 (Strasser et al. 2002; Hurt et al. 2004). In humans, the THO complex consists of THOC1 (*S*.*c*. Hpr1), THOC2 (Tho2), THOC3 (Tex1), THOC5 (Thp2), THOC6 (not present in *S*.*c*.) and THOC7 (Mft1) (Heath et al. 2016). In addition to THO, the human TREX complex contains UAP56/DEX39B (*S*.*c*. Sub2), ALYREF (*S*.*c*. Yra1) and several additional interaction partners such as SARNP (*S*.*c*. Tho1), ZC3H11A, IUF, LUZP4, POLDIP3 and CHTOP (Dufu et al. 2010; Heath et al. 2016). In addition to this high degree of conservation, the importance of TREX is also reflected by its implication in several human diseases (Heath et al. 2016).

TREX functions in transcription elongation, 3’ end processing, mRNP assembly and nuclear mRNA export (Strasser et al. 2002; Zenklusen et al. 2002; Rougemaille et al. 2008). Furthermore, TREX – in part due to the DNA-RNA helicase activity of Sub2/UAP56 (Saguez et al. 2013; Perez-Calero et al. 2020) – prevents GC-rich mRNA elements from forming so-called R-loops, DNA-RNA hybrids of the nascent mRNA with the DNA template strand that cause hyperrecombination, DNA damage and, ultimately, genome instability (Dominguez-Sanchez et al. 2011; Gomez-Gonzalez et al. 2011; Luna et al. 2019; San Martin-Alonso et al. 2021). In *S. cerevisiae*, the TREX complex is recruited to the site of transcription and the nascent mRNA by a number of interactions. THO is recruited by the C-terminal domain of RNAPII, the Prp19 complex, Mud2 and RNA (Abruzzi et al. 2004; Chanarat et al. 2011; Meinel et al. 2013; Minocha et al. 2018). THO in turn recruits Sub2, which first adopts a semi-closed conformation and upon binding to mRNA and ATP changes to a closed state (Chen 2021). Interaction of Sub2 with THO and Yra1 enhances its ATPase activity (Ren et al. 2017). Yra1 recruits the mRNA export receptor Mex67-Mtr2 to the mRNA, which exports the mRNA through the nuclear pore complex to the cytoplasm (Katahira et al. 1999; Strasser and Hurt 2000; Stutz et al. 2000; Strasser and Hurt 2001; Zenklusen et al. 2001).

The molecular mechanism and the structural basis for these different TREX activities are still not completely understood. Ren et al. determined the first structural model of the hetero-pentameric THO complex with Sub2 at a resolution of 6.0 Å by X-ray crystallography (Ren et al. 2017). This structure is based on a truncated THO* core complex, in which potentially disordered regions were removed, and revealed an elongated architecture of THO with Sub2 bound in its center (Ren et al. 2017). More recently, high resolution cryo-EM structures of human and yeast THO-Sub2 complexes with a resolution of up to 3.2 Å were reported (Puhringer et al. 2020; Schuller et al. 2020; Chen 2021; Xie et al. 2021). The first cryo-EM reconstruction of a recombinant yeast THO-Sub2 complex was published by Schuller *et al*. (Schuller et al. 2020). This structure displays a dimeric architecture, consistent with the size exclusion chromatogram. A similar cryo-EM study with heterologously expressed THO components and Sub2 was published by Chen et al. (Chen 2021). Structural insights into the interaction between THO and Gbp2 were recently obtained by combining cross-linking mass spectrometry (XL-MS) and cryo-EM (Xie et al. 2021). The obtained truncated THO*-Sub2 structure assembles in dimers as well as tetramers. The authors speculate that the tetramer is formed due to the truncations in THO* and focused on the dimeric structure for further analysis.

The arrangement of THO and Sub2 observed in the recombinant yeast structures closely resembles the cryo-EM structure of the recombinant human complex (Puhringer et al. 2020), although the human THO-UAP56 complex contains an additional protein, THOC6, which is not present in *S. cerevisiae*, and lacks a homolog of the yeast Thp2. Most recently, the structure of the endogenous human TREX complex was published (Pacheco-Fiallos et al. 2023), which overall is consistent with the structure of the recombinant complex (Puhringer et al. 2020). Both human complexes predominantly form a tetramer, although the recombinant THO-UAP56 complex also displays an octameric organization. Additionally, based on cryo-EM micrographs of TREX bound to the exon-junction complex and mRNAs a globular architecture of human TREX-mRNP complexes was suggested (Pacheco-Fiallos et al. 2023).

The existing structures of the THO-Sub2 complex obtained by EM or crystallography suggest that the organization of the complex is conserved. However, all structures obtained from recombinantly expressed proteins, depending on complex preparation, exhibit different oligomeric states ranging from dimeric to octameric (Schuller et al. 2020; Chen 2021; Xie et al. 2021).

Here, we report the structural analysis of the endogenous *S. cerevisiae* TREX complex by XL-MS, supported by negative-stain electron microscopy of the cross-linked complex, and compare it to the published recombinant yeast structures. Importantly, this is the first structural report on endogenously purified TREX complex from yeast with all nine subunits. Chemical cross-linking experiments revealed that less than 40% of the cross-links occur in the conformationally rigid regions that are resolved in the recently reported structures of recombinant THO-Sub2 complexes. Instead, many cross-links are observed in flexible regions, in line with a recent study that also reported XL-MS combined with cryo-EM for the analysis of the recombinant THO-Sub2 complex (Xie et al. 2021). Our structural analysis of the cross-linked native TREX complex using negative stain EM reveals that the endogenous *S. cerevisiae* TREX complex assembles into a dimer.

## RESULTS

### Purification of the endogenous TREX complex from *S. cerevisiae*

To obtain the *S. cerevisiae* TREX complex for structural analysis, we tagged its subunit Hpr1 endogenously with a C-terminal FLAG-TEV-Protein A tag and purified the native complex by tandem affinity purification. A representative purification of TREX is shown in Figure 1A. As expected, the THO subunits Tho2, Mft1, Thp2 and Tex1 co-purified with Hpr1 as well as the TREX components Sub2, Yra1, Gpb2 and Hrb1 (Fig. 1A, lane 1). The identity of all these components was confirmed by mass spectrometric analysis. Following purification, the complex was directly cross-linked either with BS^3^, a homobifunctional crosslinker with amine-reactive N-hydroxysulfosuccinimide esters at each end of a 6-carbon-spacer-arm (11.4 Å), or with adipic dihydrazide (ADH-d_0_/d_8_, Sigma-Aldrich) and 4-(4,6-dimethoxy-1,3,5-triazin-2-yl)-4-methylmorpholinium (DMTMM) chloride. The endogenously purified, cross-linked TREX complex was then analyzed by XL-MS and negative stain EM to gain insights into its structural arrangement.

**FIGURE 1.**
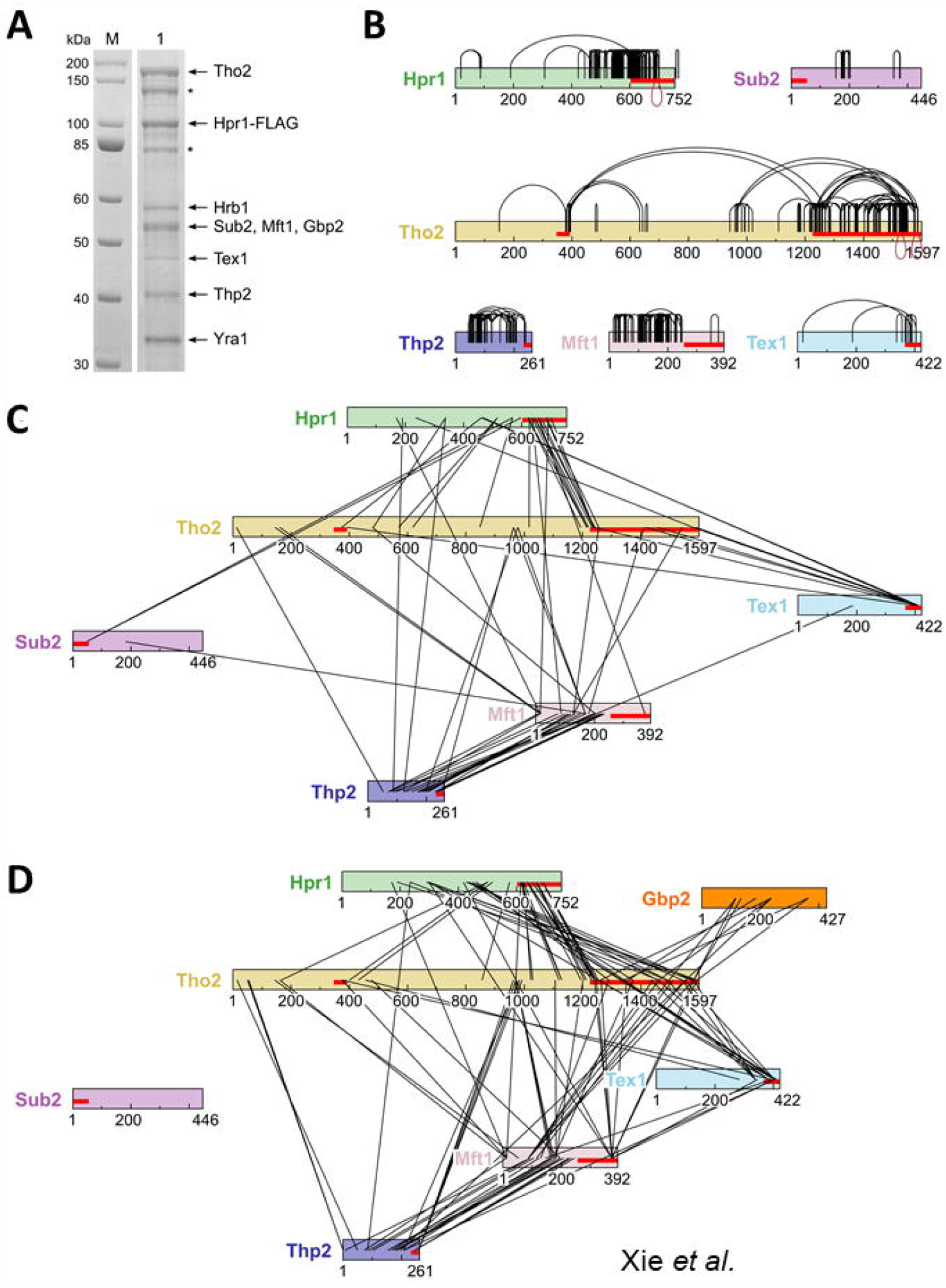
Visualization of cross-links on the TREX complex. (*A*) SDS-PAGE of the purified endogenous *S. cerevisiae* TREX complex. Subunits corresponding to Tho2, Hpr1-FLAG, Hrb1, Sub2, Mft1, Gbp2, Tex1, Thp2 and Yra1 were identified by mass spectrometry and are indicated accordingly. The asterisk indicates degradation bands of Tho2 and Hpr1. Molecular weight marker (M) bands are indicated in kDa. (*B - D*) Cross-links connecting residues within the same protein (*B*) and cross-links connecting residues of two different proteins (*C*) mapped on the sequence graphs of the corresponding proteins as black arcs and black lines, respectively. Segments longer than 25 residues that are missing in the PDB ID: 7apx structural model are shown as red lines on the sequence graph. Visualization was performed using xiNET (Combe et al. 2015) (*B*) Cross-links that unambiguously correspond to inter-molecular contacts are shown in red and facing downward. Cross-linking data from this work. (*C*) Cross-links identified in this work. (*D*) Cross-links identified by Xie et al. (Xie et al. 2021).

### Cross-linking mass spectrometry analysis of endogenously purified *S. cerevisiae* TREX complex

To investigate the molecular architecture of the endogenous yeast TREX complex, we first used XL-MS to map intra- and intermolecular contacts within the complex. Chemical cross-linking yields residue-specific amino acid contacts by forming covalent bonds between reactive side chains that are in spatial proximity (typically less than 30 to 35 Å apart). To obtain complementary information, we used two different cross-linking protocols. Crosslinking with BS^3^, which predominantly reacts with primary amines on lysines and protein N-termini, yielded many cross-links in the flexible regions of TREX components. As a second cross-linking approach, we used a combination of ADH and DMTMM chloride. This cross-linking chemistry results in two types of cross-linking products, one incorporating the dihydrazide linker, connecting carboxylic groups (aspartic or glutamic acids), the other directly coupling amines to carboxyl groups activated by the reaction with DMTMM (“zero-length” cross-links) (Leitner et al. 2014a).

Using both cross-linking chemistries, we were able to obtain more than 550 cross-linked peptide pairs, corresponding to more than 445 non-redundant site pairs on the complex subunits (THO complex and Sub2), with the majority contributed by BS^3^ and DMTMM links (Fig. 1; Supplemental Table S1). Although Yra1, Hrb2 and Gbp2 were present in the analyzed sample, no cross-links between these proteins and any of the core THO subunits were observed. This could reflect, for example, a suboptimal distribution of cross-linking sites at the respective binding interfaces, a reduced reactivity of such sites due to limited accessibility to the crosslinking reagents or the presence of these proteins at sub-stoichiometric levels. Cross-links on individual subunits were heterogeneously distributed, with the largest numbers found on the C-terminal regions of Tho2 and Hpr1, the first approximately 250 residues of Mft1 and on Thp2 (Fig. 1B). Note that cross-links between residues on the same protein may occur within one or between two copies of this protein. Cross-links connecting different subunits were most frequently observed between Tho2 and Hpr1 was well as between Mft1 and Thp2 (Fig. 1C).

### Comparison of the cross-linking data with data from recombinant yeast TREX structures

Recently, high-resolution structures of reconstituted yeast THO-Sub2 complexes have been obtained in independent studies. To assess the compatibility of our cross-linking data of the endogenous TREX complex with the reported structures, we first mapped the cross-links depicted in Fig. 1C onto the 3.4 Å cryo-EM structure from Schuller et al. (Schuller et al. 2020) [PDB ID: 7apx]. Interestingly, for less than 40% of these cross-links (171 of 445) both of the linked residues were contained in the structure; for the remaining cross-links either one or both of the connected residues are not resolved in the structure. This is especially apparent in the C-terminal part of Tho2, for which more than 350 residues are missing, and for Hpr1, for which the model lacks approximately 150 residues on the C-terminal side (Fig. 1). Unfortunately, for both proteins the rich cross-linking data in the flexible regions does not allow modeling of the full-length protein with reasonable confidence, because the flexible regions have only relatively few contacts to the more ordered regions of the proteins or to other subunits.

For the subset of cross-links that map to the structure, approximately 78% of the contacts (133 of 171) were compatible with the monomeric unit with a maximal distance of 35 Å, and an additional three cross-links matched only to subunit contacts between two monomer units in the dimer. However, 35 of the 38 remaining site pairs also do not fit the dimer structure [PDB ID: 7aqo]. The incompatible cross-links mostly reside on the subunits Thp2 and Mft1 or connect these two subunits. It should be pointed out that these cross-links were observed in multiple independent experiments and with different cross-linking chemistries. Moreover, the scores of most identifications were high so that we consider it unlikely that they correspond to false positive assignments. Therefore, we assume that these cross-links reflect different conformational or compositional states present in the native complex preparation, as we were not able to model any assembly state (monomeric, dimeric or tetrameric) that would be compatible will all cross-linking data.

We furthermore compared our cross-linking data set (Fig. 1C) to the cross-linking results of Xie *et al*. (Fig. 1D). Xie *et al*. also used XL-MS to confirm intra- and inter-molecular contacts in a reconstituted yeast THO-Sub2-Gbp2 complex, in their case using the amine-reactive disuccinimidyl suberate (DSS), a non-sulfonated analog of BS^3^, and the zero-length coupling reagent N-(3-Dimethylaminopropyl)-N′-ethylcarbodiimide (EDC) (Xie et al. 2021). Even more so than for our data set, a large fraction (243 of 314, approximately 75%) of the DSS/EDC links could be mapped to the structure determined by Schuller et al. (2020). In contrast to our data from the natively purified endogenous complex, the amino acid contacts that were covered in the structure were compatible with the 35 Å threshold. The higher fraction of compatible cross-links probably reflects the higher structural homogeneity for the recombinant complex. Figure 2 compares the inter-protein cross-links observed in both studies and shows that the most frequently covered subunit-subunit contacts are quite similar.

**FIGURE 2.**
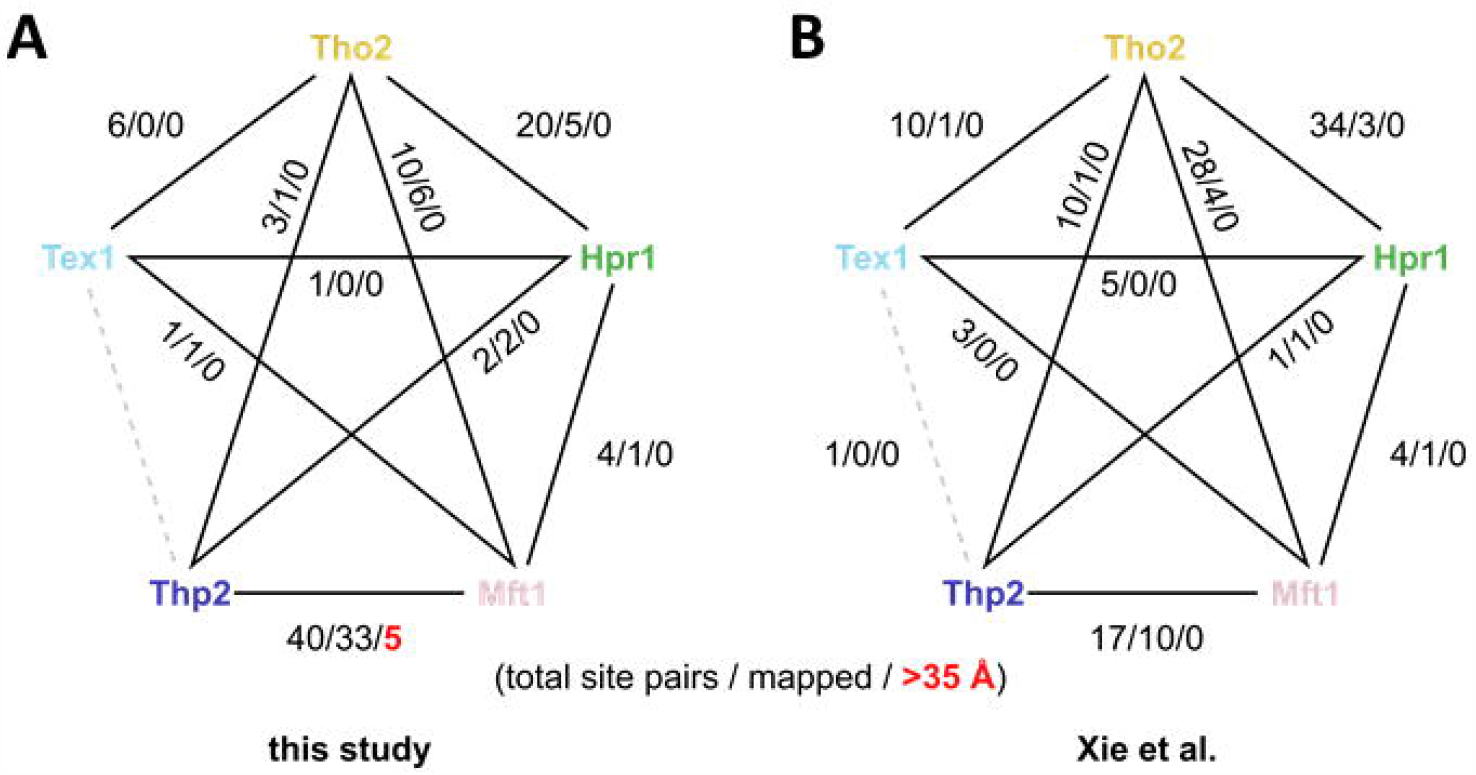
Comparison of inter-subunit cross-links. (*A*) observed in our work, excluding contacts with Sub2, and (*B*) from cross-linking of the recombinant complex by Xie *et al*. (Xie et al. 2021). Shown are the numbers of non-redundant site pairs observed in total for each subunit pair, the number of contacts that could be mapped onto PDB ID: 7apx, and (in red) those that exceed the distance of 35 Å. Proteins are color-coded as in Figure 1.

### Structure determination of the cross-linked TREX complex by negative stain EM

Since a majority (over 60%) of the cross-links observed in our XL-MS analysis are located in regions that are not resolved in the recently published THO-Sub2 structures, e.g. (Schuller et al. 2020), we next examined whether formation of the endogenous TREX complex differs from that of the recombinant THO-Sub2 complex. We were especially interested in the preferred oligomeric state of the endogenous TREX complex purified from its native environment, since the XL-MS data suggests different conformational or compositional states present in the sample. We therefore subjected freshly BS^3^ cross-linked complex to uranyl acetate negative stain (UANS) electron microscopy (EM) and single-particle analysis (Supplemental Fig. S1). The sample showed a heterogeneous complex formation on the negative stain grid (Fig. 3A); however, distinct dimer-like particles similar to previously published 2D classes were observed (Fig. 3B) (Pena et al. 2012; Schuller et al. 2020; Xie et al. 2021). To estimate the fraction of dimer-like particles in the UANS dataset, we compared reference-free 2D class averages to the published dimer map of the yeast THO-Sub2 complex by Schuller et al. using image cross-correlation techniques (Supplemental Fig. S1A). When opting for a cross-correlation coefficient cut-off value that includes distinct dimer views into the dataset, the dimer-correlated fraction in the UANS dataset is 50%, with another 27% at coefficients between 0.897 and 0.85. We thus conclude that the majority of particles in the dataset have a dimer-like appearance. Subsequently, the dimer fraction was used to generate a 3D reconstruction without any symmetry applied, yielding an overall resolution of 26 Å (Fig. 3C). Comparison of the previously published yeast THO-Sub2 dimer (Schuller et al. 2020) with our 3D reconstruction or the reference free class averages reveals a strong similarity between the two complexes (Fig. 3D and E, Supplemental Fig. S1A). At the resolution of the UANS dataset, the endogenously purified *S. cerevisiae* TREX sample resembles the high-resolution structure of the *in vitro* assembled complex in terms of overall shape and order with small differences in domain orientations (Fig. 3E and F).

**FIGURE 3.**
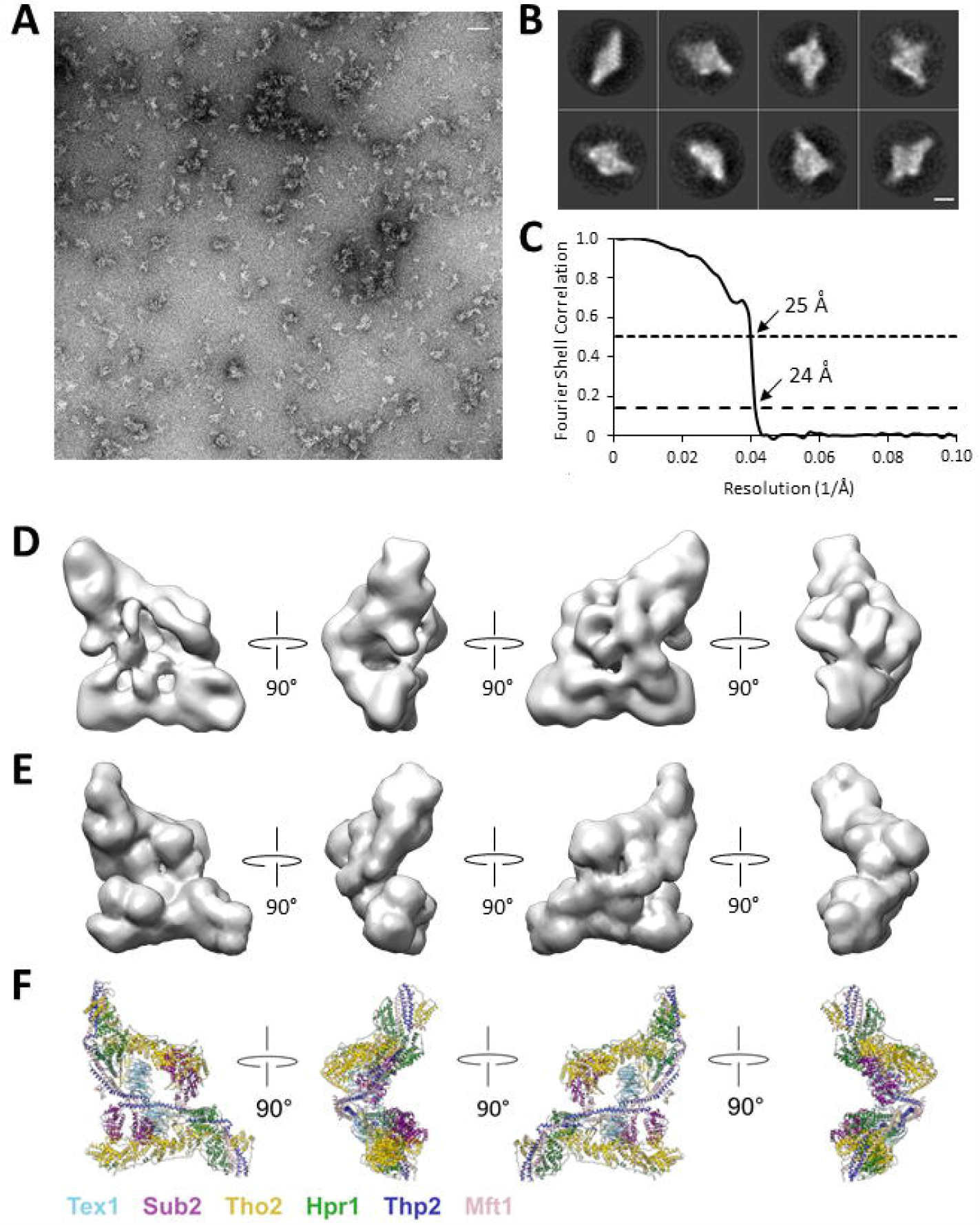
Negative stain EM structure of the endogenous TREX complex from *S. cerevisiae*. (*A*) A representative negative stain micrograph of the BS^3^ cross-linked *S. cerevisiae* TREX complex. Scale bar represents 500 Å. (*B*) Representative reference-free 2D class averages of the *S*.*c*. TREX complex. Scale bar represents 100 Å. (*C*) Fourier Shell Correlation (FSC) plot for the negative stain reconstruction of the *S*.*c*. TREX complex. The resolution at FSC=0.143 (dashed line) and FSC=0.5 (dotted line) are indicated. (*D*) Three-dimensional reconstruction of the *S*.*c*. TREX complex resolved to 26 Å. (*E*) Representative views of the Tho-Sub2 dimer structure (PDB ID: 7aqo) in surface representation lowpass-filtered to 25 Å. (*F*) Color-coded THO-Sub2 dimer PDB model depicted in the same views as in (*E*). Proteins are color-coded as in Figure 1.

## DISCUSSION

The TREX complex is essential for gene expression and plays crucial roles in transcription elongation, 3’ end processing, mRNP assembly, nuclear export and genome stability. Recently, several high-resolution structures of the human and *S. cerevisiae* THO-Sub2 complex were published that elucidate TREX function from a structural perspective (Puhringer et al. 2020; Schuller et al. 2020; Chen 2021; Xie et al. 2021). In most of these works, protein subunit purification and *in vitro* complex reconstitution were optimized to yield highly homogeneous complexes, e.g., by truncation of flexible regions and a combination of prokaryotic and eukaryotic expression systems. However, although the resulting structural models are highly similar with respect to the monomeric complexes, they exhibit different oligomeric states, raising the concern that the recombinant structures may not reflect complex organization in the native cellular environment of yeast. Overall, our XL data of the native TREX complex of *S. cerevisiae* supports the structures of the recombinant TREX subcomplex and thus confirms the validity of the recent high-resolution structures for the interpretation of *in vivo* functional data.

Interestingly, the majority of our cross-linked residues (60%) are located in flexible regions that are not resolved in the recently reported cryo-EM structures of the recombinant THO-Sub2/UAP56 complexes (Puhringer et al. 2020; Schuller et al. 2020; Chen 2021; Xie et al. 2021), similar to a recent study that also employed XL-MS for the analysis of the recombinant THO-Sub2 complex (Xie et al. 2021). In particular, this concerned the C-terminal tails of Tho2, Hpr1, and – to a lesser extent – Tex1 (Fig. 1). The observed cross-linking pattern is not untypical for flexible regions, however, according to our experience, the strong preference for these regions of the complex is unusual. Furthermore, we were unable to find conditions that resulted in a more balanced distribution of cross-links over the entire surface. XL-MS is frequently used to provide additional structural information on regions that are not accessible to classical structural biology techniques, as the distance restraints can be used as input for integrative/hybrid modelling approaches (Braitbard et al. 2019; Rout and Sali 2019; Koukos and Bonvin 2020). However, in this case, most of the intra-molecular cross-link contacts connect regions that are relatively close to each other in the primary sequence (i.e., within the flexible regions), and with relatively few contacts to more ordered regions of the complex, so that a restraint-based modelling approach is unlikely to be successful.

Among the cross-links covered by reported structures of the recombinant complex, close to 80% are compatible with a maximal distance of 35 Å between the involved residues. The remaining 20% could either be the result of cross-linking induced artefacts, or they could reflect other subunit arrangements in the natively purified complex. However, we argue against the former explanation for several reasons. First, cross-linking conditions were optimized to avoid excessive cross-linking. For example, the low reagent concentrations for the ADH + DMTMM reaction resulted in relatively few cross-links incorporating the ADH linker, as expected from recent observations (Mohammadi et al. 2021), and favored DMTMM-induced zero-length cross-links. Xie et al. used a DSS cross-linker concentration identical to our BS^3^ conditions (0.5 mM), and their results show a better compatibility with the model from Schuller et al. (2020). Given that DSS and BS^3^ are comparable in their reactivity, this also points to a limited influence of cross-linking conditions, and rather argues for a conformational difference of at least a subset of the respective purified complexes. Such differences in composition or conformation can be enhanced or even distorted in XL-MS experiments since contacts from lowly populated states may still result in high-confidence identifications if such cross-linked peptide pairs ionize and fragment well.

The negative stain EM structure of the cross-linked endogenous *S. cerevisiae* TREX complex presented here in overall appearance corresponds reasonably well to the published cryo-EM structures of the recombinant THO-Sub2 complex (Schuller et al. 2020). Our endogenous TREX complex is predominantly dimeric and supports the findings for the recombinant THO-Sub2 complex, although the endogenously purified complex we model here additionally contains Hrb1, Gbp2, Yra1 and the C-terminal 42 residues of Tex1 that were truncated in the recombinantly purified complexes. Small differences in domain orientation and extra densities between the native and recombinant structures are most likely due to the additional proteins, which unfortunately cannot be located without doubt at this resolution. However, the additional proteins present in our purification do not seem to cause the complex to adopt major alternative conformations.

Pacheco-Fiallos *et al*. (2023) showed that the endogenous human TREX complex is structurally consistent with the recombinant human THO-UAP56 complex (Puhringer et al. 2020). The only structure of endogenous, fully assembled yeast TREX complex so far shows an elongated, mainly monomeric assembly in negative stain EM (Pena et al. 2012), which only partially fits our dimeric reconstruction as the dimer arrangement described could not be observed in our dataset. Intriguingly, one protomer of the recombinant THO-Sub2 dimer complex (Schuller et al. 2020) fits well into our EM density, while the second protomer leaves some unfilled density. This could hint towards additional interactions with TREX subunits identified in our endogenously purified complex by MS (Yra1, Gbp2, Hrb1) taht are not present in the structure of the heterologously expressed THO-Sub2 (Schuller et al. 2020; Xie et al. 2021). However, the unequal density distribution could also be due to differing stoichiometries of TREX components. Nevertheless, apart from these minor differences, we could not identify a major alternative conformation that would explain the 35 violating cross-links of our XL-MS analysis that were not compatible with the dimer structure from Schuller et al. (2020) or other assembly states like monomer or tetramer. We thus suggest that these cross-links could derive either from aggregated protein or from assembly intermediates of TREX complexes that are only present in the natively purified sample.

In summary, based on our XL-MS and negative stain EM analyses, the structure of the native TREX complex of *S. cerevisiae* corresponds to the structures determined for the recombinant THO-Sub2 complex confirming the validity of the recent high-resolution structures for the interpretation of *in vivo* functional data.

## MATERIALS AND METHODS

### Purification of the native yeast TREX complex

A sequence coding for a FLAG-TEV-2xProtein-A (FTpA) tandem affinity tag was inserted C-terminally into the endogenous *HPR1* locus into the *S. cerevisiae* strain RS453 (MATa) using standard yeast genetic techniques. Yeast was grown in 30 l YPD to OD_600_ = 3.5, harvested by centrifugation and frozen in liquid nitrogen. Frozen yeast pellets were lysed with a cryo-mill (SPEX) and stored at -80°C. The grindate was resuspended in buffer A (50 mM HEPES, pH 7.5; 200 mM KCl; 1.5 mM MgCl_2_; 0.15% IGEPAL) containing protease inhibitor mix (2 µM pepstatin A, 0.6 µM leupeptin, 1 mM PMSF and 2 mM benzamidine). The lysate was pre-cleared at 4,200 g for 10 min at 4°C, followed by ultracentrifugation of the supernatant at 132,600 g for 1 h at 4°C. The clear supernatant was transferred to four 50 ml conical centrifuge tubes. Each tube was rotated with 600 µl IgG beads (Cytiva) for 16-18 h at 4°C. Beads were separated from the lysate by centrifugation at 700 g for 3 min at 4°C and washed twice with buffer A containing 1 mM DTT. For elution, beads were rotated with TEV protease in buffer A for 1 h at 16°C. Following centrifugation, the resulting supernatant (TEV eluate) was rotated with 450 µl FLAG beads (Sigma-Aldrich) in buffer B (20 mM HEPES, pH 8.4; 150 mM KCl) or buffer C (20 mM HEPES, pH 7.0; 150 mM KCl) for 1 h at 4°C. Beads were washed once with buffer B or C containing 0.15% IGEPAL. For elution, the sample was rotated with 100 µg/ml FLAG peptide (Sigma-Aldrich) for 15 min at room temperature. The purified *S*.*c*. TREX complex was concentrated and subsequently cross-linked with either bis(sulfosuccinimidyl)suberate (BS^3^-d_0_/d_12_, Creative Molecules) in buffer B or with adipic dihydrazide (ADH-d_0_/d_8_, Sigma-Aldrich) and 4-(4,6-dimethoxy-1,3,5-triazin-2-yl)-4-methylmorpholinium (DMTMM) chloride (Sigma-Aldrich) in buffer C for 1 h at 37°C and 500 rpm using a ThermoMixer (Eppendorf). The BS^3^ cross-linking reaction was stopped by incubation with 50 mM ammonium bicarbonate (NH_4_HCO_3_) for 20 min at 37°C at 500 rpm in a ThermoMixer. The ADH/DMTMM cross-link reaction was stopped by desalting with a Zeba™ Spin Desalting Column (ThermoFisher Scientific). The cross-linked samples were flash frozen in liquid nitrogen and stored at -80°C until further processing. For the XL-MS analyses, 40 µg purified *S*.*c*. TREX complex each were cross-linked in three independent experiments with (1) 0.5 mM BS^3^, (2) 1 mM ADH and 1 mM DMTMM or (3) 2.5 mM ADH and 2.5 mM DMTMM. For EM analysis, 20 µg *S*.*c*. TREX sample was cross-linked with BS^3^.

### Sample processing for mass spectrometry

After quenching or gel filtration, cross-linked samples were evaporated to dryness in a vacuum centrifuge and processed essentially as described previously (Leitner et al. 2014b). In brief, disulfide bonds were reduced with tris(2-carboxyethyl)phosphine and free thiol groups in cysteines were alkylated with iodoacetamide. Proteins were digested in a two-step procedure, first with endoproteinase Lys-C (Wako; 1:100, 37°C for 2-3 h), then with trypsin (Promega; 1:50, 37°C overnight). Samples were purified by solid-phase extraction (SepPak tC18 cartidges, Waters) and fractionated by peptide-level size-exclusion chromatography (SEC; Superdex Peptide PC 3.2/300, GE) on an ÄKTA micro FPLC system (Leitner et al. 2012).

### Liquid chromatography-tandem mass spectrometry

SEC fractions and additionally some unfractionated samples were analyzed by nanoflow LC-MS/MS. The BS^3^ sample was initially analyzed on an Orbitrap Elite system and later reanalyzed on an Orbitrap Fusion Lumos instrument (both ThermoFisher Scientific), while the ADH/DMTMM samples were only analyzed on the latter instrument.

The Orbitrap Elite instrument was connected to an Easy nLC-1000 HPLC system (ThermoFisher Scientific) with a 150 mm × 75 µm Acclaim PepMap RSLC C_18_ column (2 µm particle size, ThermoFisher Scientific) and mobile phases A = water/acetonitrile/formic acid (98:2:0.15, v/v/v) and B = acetonitrile/water/formic acid (98:2:0.15, v/v/v). Peptides were separated at a flow rate of 300 nL/min with a gradient of 9 to 35% B in 60 min. The mass spectrometer was operated in data-dependent acquisition mode with detection of precursor ions in the Orbitrap at 120000 resolution and detection of fragment ions in the linear ion trap at normal resolution. For each sequencing cycle, the 10 most abundant precursor ions (top n mode) were selected for sequencing if they had a charge state of +3 or higher and were fragmented at 35% normalized collision energy. Dynamic exclusion was activated for 30 s after one sequencing event.

The Orbitrap Fusion Lumos mass spectrometer was connected to an Easy nLC-1200 HPLC instrument (ThermoFisher Scientific) with a 250 mm × 75 µm Acclaim PepMap RSLC C_18_ column (2 µm particle size, ThermoFisher Scientific) and mobile phases A = water/acetonitrile/formic acid (98:2:0.15, v/v/v) and B = acetonitrile/water/formic acid (80:20:0.15, v/v/v). Peptides were separated at a flow rate of 300 nL/min with a gradient of 11 to 40% B in 60 min. The mass spectrometer was operated in data-dependent acquisition mode with detection of precursor ions in the Orbitrap at 120,000 resolution and detection of fragment ions in the linear ion trap at rapid resolution (low-resolution MS/MS) or in the Orbitrap at 30,000 resolution (high-resolution MS/MS). For each sequencing cycle, the most abundant precursor ions were selected for sequencing with a cycle time of 3 s (top speed mode) if they had a charge state of +3 or higher and were fragmented at 35% normalized collision energy. Dynamic exclusion was activated for 30 s after one sequencing event.

### Identification of cross-linked peptides

MS/MS data acquired in the native Thermo .raw format were converted into the mzXML format by msconvert/ProteoWizard (Chambers et al. 2012) and searched using xQuest (Walzthoeni et al. 2012), version 2.1.4, against a custom database of target proteins (subunits of the TREX complex) and identified contaminants. The false discovery rate (FDR) was controlled by a target/decoy search using reversed and shuffled sequences as decoys. xQuest search parameters included: enzyme = trypsin, maximum number of missed cleavages = 2, fixed modifications = carbamidomethylation on Cys, initial MS1 error tolerance = ±15 ppm, MS2 error tolerance = 0.2 Da for “common” fragment ions and 0.3 Da for “xlink” fragment ions (ion trap) or = ±20 ppm (Orbitrap), respectively. Cross-linking specificity was set to K and N terminus for BS^3^, D or E for ADH, and K with D or E for DMTMM. Primary search results (target and decoy hits) were then further filtered according to the experimentally observed mass accuracy, a TIC value of >0.1 (for BS^3^ or ADH) or >0.15 (for DMTMM) and a deltaS value of <0.9. All remaining candidate hits were further curated and only assignments for which a minimum of four bond cleavages overall or three consecutive bond cleavages were assigned for each peptide were retained. The FDR was adjusted to 5% at the cross-linked peptide pair level after the manual curation steps.

### Structural validation of cross-links

Cross-linking data from our own experiments or retrieved from published work (Xie et al. 2021) were mapped on the structure of the THO-Sub2 complex from Schuller et al. (Schuller et al. 2020) [monomer: PDB ID: 7apx, dimer: PDB: 7aqo] using PyXlinkViewer (Schiffrin et al. 2020). An upper bound Euclidean distance of 35 Å (C_α_-C_α_) was selected for “compatible” cross-links. Visualization of cross-links for Figure 2 was performed using xiNET (Combe et al. 2015).

### Negative stain electron microscopy, data collection and data processing

3.5 µl of BS^3^ cross-linked *S*.*c*. TREX complex (c = 60 µg/ml) were applied to a freshly glow-discharged carbon-coated copper mesh grid and stained two times with 2% uranyl acetate. Negative stain EM images were collected on a Talos F200C TEM microscope (Thermo Fisher) operated at 200 kV equipped with a Ceta 16M camera (Thermo Fisher). 599 micrographs were recorded at a magnification of 54 kx at a pixel size of 2.54 Å using EPU. The defocus range was set from 1.5 to 2.5 µm.

Image processing and 3D reconstruction was performed using Relion-3.0 (Zivanov et al. 2018). Contrast transfer functions were determined using CTFFIND4 (Rohou and Grigorieff 2015). All refinements used gold standard Fourier shell correlation (FSC) calculations and reported resolutions are based on the FSC = 0.143 criterion. 2D classification of 630 manually picked particles generated templates for semi-automated particle selection. The dataset of 88,543 picked particles was subjected to several rounds of unsupervised 2D classification to remove bad particles resulting in a final dataset of 42,784 particles. The 2D class averages of the 42,784 particle set and a Molmap of PDB ID: 7aqo lowpass-filtered to 25 Å were subjected to projection matching in IMAGIC (van Heel and Keegstra 1981) to analyse dimer representation in the dataset. Only 2D class averages with a cross-correlation coefficient larger 0.897 were selected resulting in a final particle set of 22,024 particles. Selected particles were used for a 3D refinement in CryoSPARC v4 (Punjani et al. 2017). The EM map of the endogenous *S. cerevisiae* TREX complex was deposited under accession code EMD-17464.

## DATA DEPOSITION

All XL-MS data have been deposited to the PRIDE repository (45) (PXD025843), and all identified cross-linked peptide pairs are listed in Supplemental Table S1. The EM map of the endogenous *S. cerevisiae* TREX complex was deposited under accession code EMD-17464.

## SUPPLEMENTAL MATERIAL

Supplemental Material is available for this article.

## ACKNOWLEDGEMENTS

We thank Manuel Koschitza for initial experiments for this project, Birte Keil for purification experiments of THO truncation mutants, Cornelia Kilchert and Vera Bettenworth for critically reading the manuscript. A.L. would like to thank Ruedi Aebersold and Paola Picotti for access to laboratory infrastructure and instrumentation. EM data were recorded on the Talos F200 supported by the DFG under INST 336/148-1 FUGG. We thank Petra Wendler for helpful discussions and for providing microscope data. C.R. thanks Andreas Heimann for IT support. This work was supported by the European Union [European Research Council Consolidator Grant, grant number 772049 to K.S. and Grant ULTRA-DD, grant number FP7-JTI 115766] and ETH Zurich [Scientific Equipment Grants].

